# Fibrinogen and Complement Factor H are promising CSF protein biomarker(s) for Parkinson’s disease with cognitive impairment- A Proteomics and ELISA based study

**DOI:** 10.1101/2021.03.19.436097

**Authors:** Aditi Naskar, Albert Stezin, Arpitha Dharmappa, Shantala Hegde, Mariamma Philip, Nitish Kamble, Jitender Saini, K Sandhya, Utpal Tatu, Ravi Yadav, Pramod Kumar Pal, Phalguni Anand Alladi

## Abstract

Cognitive impairment is a debilitating non-motor symptom of Parkinson’s disease (PD). The diagnosis of PD with cognitive impairment (PDCI) is essentially through clinical and neuropsychological examinations. There is an emerging need to identify biomarker(s) to foresee cognitive decline in PD patients, at an early stage. We performed label-free unbiased nontargeted proteomics (Q-TOF LC/MS-MS) in CSF of non-neurological control (NNC); PDCI; PD and normal pressure hydrocephalus (NPH), followed by targeted ELISA for validation. The diagnosis was confirmed by neuropsychological and MRI assessments prior to CSF collection. Of the 282 proteins identified by mass spectrometry, 42 were differentially altered in PD, PDCI and NPH. Further, 28 proteins were altered in PDCI and 25 in NPH. An interesting overlap of certain proteins was noted both in PDCI and NPH. Five significantly upregulated proteins in PDCI were fibrinogen, gelsolin, complement factor-H, apolipoprotein A-IV and apolipoprotein A-I. Whereas carnosine dipeptidase 1, carboxypeptidase E, dickkpof 3 and secretogranin 3 precursor proteins were down-regulated. NPH also had few uniquely altered proteins viz. insulin-like growth factor-binding protein, ceruloplasmin, α-1 antitrypsin, VGF nerve growth factor, neural cell adhesion molecule L1 like protein. Interestingly, the ELISA-derived protein concentrations correlated well with the neuropsychological scores of certain cognitive domains. Executive function was affected both in PDCI and NPH. In PD, Wisconsin card sorting test (WCST) percentile correlated positively with ApoA-IV and negatively with the ratio of ApoAI: ApoA-IV. Thus assessment of a battery of proteins like fibrinogen-α-chain, CFAH and ApoAI: ApoA-IV ratio alongside neuropsychological could be reliable biomarkers to distinguish PDCI and NPH.

## Background

Currently, the understanding of Parkinson’s disease (PD) has shifted from being a completely motor disorder to a multisystem disorder that is accompanied by non-motor symptoms. Cognitive impairment is the most widespread and debilitating non-motor symptom of PD. It ranges from tenuous cognitive decline in mild cognitive impairment (MCI) to severe cognitive impairment resulting in dementia i.e. PD with dementia (PDD). The term mild cognitive impairment (MCI) is often used in Alzheimer’s disease (AD) as a transitional state between normal cognition and dementia [1] and has been recently introduced in the investigations related to PD [2]. Dementia is a major late-stage cognitive complication of PD. As per the Sydney multicenter longitudinal study (20 years follow-up), about 20-33% of the PD patients suffered from mild cognitive impairment (MCI) at the time of diagnosis and 80% developed dementia over the course of the disease [3]. In India, a hospital based 7 years longitudinal study showed cognitive impairment in 49% of PD patients [4]. The disease phenotype of PDD often resembles that of dementia with Lewy bodies. Therefore, the symptoms of dementia must appear at least one year after the motor impairment, for it to be termed as PDD [5]. If dementia predates PD or is diagnosed within one year of motor symptoms, the patients are categorized as DLB [5]. The pathology underlying cognitive impairment in PDD is heterogeneous and includes Lewy bodies (LB), neurofibrillary tangles, senile plaques, AD like pathology. The presence of cortical LB is tightly associated with cognitive impairment [6,].

The two different phenotypes of Parkinson’s disease with cognitive impairment (PDCI) involve fronto-striatal and posterio-cortical regions [7]. The underlying cause of fronto-striatal phenotypes are loss of dopaminergic neurons and characterized by executive dysfunction such as deficits in planning, working memory, attentional set-shifting, as well as impaired memory recall, reinforcement learning and inhibition [8]. PD patients with fronto-striatal phenotypes are less likely to progress to dementia. The posterior-cortical phenotypes involve additional neurotransmitter deficits, besides dopamine. In this phenotype visual memory deficits dominate, while deficits in the language and memory co-exist. PD patients with posterior cortical phenotypes are prone to dementia. However, little is known about the pathophysiology of cognitive decline in PD patients.

Normal pressure hydrocephalus (NPH) is a non-neurodegenerative disease characterized by a gamut of symptoms viz. gait disturbance, urinary incontinence and dementia. It is associated with ventricular enlargement with normal intracranial pressure and impaired cerebrospinal fluid (CSF) absorption. The cognitive deficits in NPH include decline in memory, attention and executive functions and is predominated by frontal and subcortical dysfunctions [9]. In view of overlap between the cognitive dysfunctions in PDCI and NPH, we included the latter group as an internal disease control for comparison.

Hitherto, the prediction of the risk and time of onset of cognitive impairments in PD (PDCI) remains clinically daunting due its heterogeneous pathology and overlap with other PD mimics. The final confirmation is possible only at autopsy. This highlights the unmet need to identify biomarker(s) to predict PDCI at an early stage. A biomarker is “a characteristic that is objectively measured and evaluated as an indicator of normal biological processes, pathogenic processes, or pharmacologic responses to a therapeutic intervention.” (Biomarkers Definitions Working Group: NIH). Most biomarker studies conducted till date for PDCI, focused on targeted CSF markers of AD i.e. Aβ42, T-tau & P-tau; whereas only a handful used non-targeted proteomics [10]. The proteins like ApoA-I and clusterin identified by non-targeted proteomics did not match those reported in PD pathogenesis [10]. Similarly, inconsistencies plague the findings of total CSF α-synuclein in different studies, since a set of studies showed reduced levels in PDD [11, 12] while others showed an increase [13]. A thin overlap between AD and PDCI pathologies pushes a likelihood of independent pathways in PDCI as against AD.

We initiated the study by estimating α-synuclein levels in PDCI-CSF by ELISA, to verify findings of the studies reported elsewhere. In view of inconclusive results, we used unbiased label-free LC-MS/MS mass spectrometry. We further validated the significantly upregulated proteins by ELISA. The CSF samples of non-neurological controls (NNC), PD, PDCI and NPH were investigated simultaneously.

## Materials & Methods

### Subject recruitment and clinical evaluation

The study was approved by the institutional human ethics committee [IEC No. NIMHANS/1st IEC (BS & NS DIV.)/2016]. All subjects/patients were recruited after obtaining written informed consent and documentation of their demographic details (Table. 1). PD, PDCI and NPH Patients were recruited over a period of 3 years (2016-2018) from the movement disorders outpatient clinic at the department of Neurology, National Institute of Mental Health & Neuro Sciences, Bengaluru, India. Idiopathic PD was diagnosed based on the UK Parkinson’s Disease Society Brain Bank criteria (UKPDS, Hughes et al. 1993). The clinical details such as, age at onset of motor symptoms, disease duration, UPDRS-III, OFF-ON state scores were documented for each patient. NPH was diagnosed based on the international iNPH and the Japanese iNPH guidelines [14] by movement disorder specialists (AS, NK, RY, & PKP). CSF samples donated by otherwise healthy individuals, not suffering from any neurological or movement disorders and undergoing spinal anesthesia, were considered as NNC. Their CSF was collected by an anesthetist (SK) at the department of Plastic Surgery, at Bangalore Medical College Research Institute. All patients underwent neurological and cognitive examinations followed by MRI-brain (JS) and routine laboratory investigations. The patients and NNCs were matched for age and socio-economic status.

**Table 1:** Demographic details of the patients enrolled in the study.

| **Characteristics** | **NNC** | **PD** | **PDCI** | **NPH** |
| --- | --- | --- | --- | --- |
| **Total No. of subjects/**<br>**patients** | 13 | 6 | 11 | 12 |
| **Sex** | M=9, F=4 | M=5, F=1 | M=11, F=0 | M=11, F=1 |
| **Median age**<br>**(Years)** | 54<br>(45-71) | 60<br>(53-67) | 65<br>(52-73) | 62<br>(50-73) |
| **Mean disease**<br>**duration**<br>**(Years)** | NA | 6 | 3 | 4 |
| **Median**<br>**MOCA score** | 22<br>(18-28) | 24.5<br>(24-28) | 20<br>(8-24) | 19.5<br>(7-30) |
| **Median**<br>**UPDRS score**<br>**'ON' state** | NA | 25.25<br>(18-26.5) | 40 (12.5-64) | 20<br>(7-44) |
| **Median**<br>**UPDRS score**<br>**'OFF' state** | NA | 30<br>(18-41) | 37<br>(37-77) | 27.5<br>(12-19) |

### Neuropsychological tests

We used Montreal cognitive assessment (MOCA) test, with a cut-off score of 26 for neuropsychological screening of patients (n=60). The patients with scores <26 underwent detailed assessment (AD, AN, SH), using selected tests from the NIMHANS neuropsychology battery. The raw scores were compared with age, education, and gender-based norms [15]. A score <15^th^ percentile was considered as a neuropsychological deficit [16]. Five cognitive domains, viz. attention, language, learning, executive function, and visual memory function were assessed for each patient. Two types of cognitive impairment manifestations were considered to identify PD-MCI and PDD. The PD-MCI was diagnosed based on MDS task force criteria, [17] with abnormalities in two tests under a single cognitive domain or abnormalities in at least one test in two cognitive domains whereas; abnormalities in more than two domains were a pre-requisite for PDD. In view of the small sample size the PD-MCI and PDD patients were grouped together as PDCI. The details of the neuropsychological tests are provided in (Table 2).

**Table 2:**
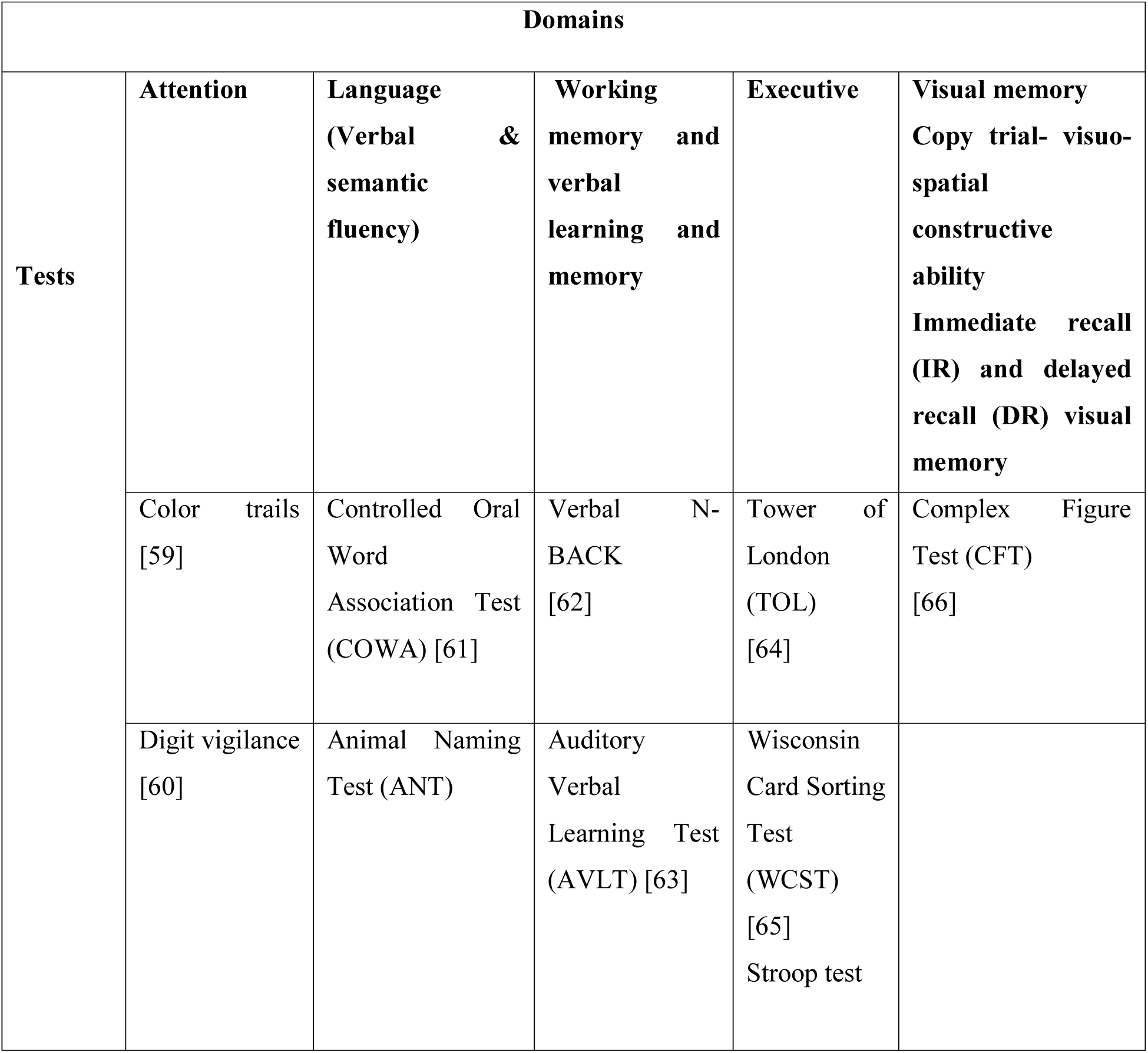
Neuropsychological tests studied under each domain

| Domains |  |  |  |  |  |
| --- | --- | --- | --- | --- | --- |
| Tests | Attention | Language<br>(Verbal & semantic fluency) | Working<br>memory and verbal learning and memory | Executive | Visual memory<br>Copy trial- visuo-spatial<br>constructive ability<br>Immediate recall (IR) and delayed recall (DR) visual memory |
|  | Color trails [59] | Controlled Oral Word Association Test (COWA) [61] | Verbal N- BACK [62] | Tower of London (TOL) [64] | Complex Figure Test (CFT) [66] |
|  | Digit vigilance [60] | Animal Naming Test (ANT) | Auditory Verbal Learning Test (AVLT) [63] | Wisconsin Card Sorting Test (WCST) [65]<br>Stroop test |  |

### Neuroimaging methods

Magnetic resonance imaging (MRI) being a supportive test for NPH, this modality was used to screen PD, PDCI and NPH patients. For the routine diagnostic MRI, T2 weighted imaging (T2 WI) was performed in all three planes (axial, coronal and sagittal) in addition to fluid attenuation inversion recovery (FLAIR), T1 weighted imaging (T1W), diffusion weighted imaging (DWI) and susceptibility weighted imaging (SWI) in the axial plane. Representative images are provided in supplementary data (S1).

### Sample collection and handling

The CSF samples were collected at room temperature by standard lumbar puncture with aseptic precautions and stored in polypropylene tubes. Within half an hour, the samples were centrifuged at 10,000 rpm (Eppendorf USA) for 10 min at 4°C to pellet out cell debris and visually inspected for blood contamination. Blood free samples were stored at −80°C for analysis.

### Mass spectrometry

a. **Sample filtration, protein estimation and preparation**

All the samples were processed and analyzed individually (NNC, n=7; PD, n=4; PDCI, n= 9; NHP, n=10). We passed 500 µl of CSF through Amicon filter (3 Kda cut-off) and determined protein concentration for 300 µl of the supernatant by Bradford assay. 50 µg of protein (approximately 225 µl) was processed further. Proteins were denatured by heat at 60°C for 10 min. Thereafter the samples were alkylated with 10 µl of 100 mM dithiothreitol (DTT) at 60°C for 25 min to break the disulfide bonds. The alkylated samples were further reduced by 10 µl of 200 mM iodoacetamide (IAA) at room temperature for ∼45 min in dark. Thereafter, the samples were digested using trypsin (overnight at 37°C, ratio 1:50), quenched using 10 µl of formic acid and centrifuged for 5 min at 13000 rpm. Post-centrifugation, the supernatant was vacuum dried using a Speed Vac (Eppendorf, USA) and reconstituted with a solution of 0.1% formic acid (FA) and 3% acetonitrile (ACN) in mass spectrometry grade water [18]

### LC-MS/MS analysis

Protein detection and identification was performed using Agilent Technologies 6545XT Advance Bio LC/Q-TOF equipped with Agilent 1290 Infinity II LC System. Data acquisition was controlled by Mass-Hunter workstation data acquisition software (B.08.01, Agilent). The mobile phases used for liquid chromatography included buffers A (Milliq water/0.1% FA) and B (ACN/ 0.1% FA). We used Agilent Advance Bio Peptide Map (2.1 ×150 mm, 2.7µ) column for chromatography. The peptides were separated by a 125 min gradient at flow rate of 0.4 ml/min. The gradient was as follows: from 2 to 35% buffer B for 100 min, then from 35% to 90% buffer B for 110 min. After rinsing with 90% B for 5 min, the column was equilibrated by 98% buffer A for 10 min. The Advance Bio Q-TOF was operated under capillary voltage of 3500V. The drying gas flow rate and temperature was set at 13 ml/min and 325°C, respectively. The Q-TOF was run in a positive mode and MS scans were run over a range of m/z 300–1700 at 8 spectra/ second. Precursor ions were selected for auto MS/ MS at an absolute threshold of 1000 and with maximum of 10 precursors per cycle, and active exclusion set at 1 spectra and released after 0.15 min reference mass correction was performed by using mass 922.0097 (±10 ppm).

### Mass Spectrometry Data analysis

The data obtained were subjected to MaxQuant Andromeda software (version 1.6.14.0.) [19] mediated search against the uniport FASTA dataset of human proteome. The Perseus software version 1.6.14.0 was used to obtain the quantitative data for all peptides by selecting the peak intensities across the whole set of protein intensity measurements. Intensity based ‘label free quantitation (LFQ)’ parameter was applied, values were imported and transformed into logarithmic scale with base 2. Based on the normal distribution, the missing values were imputed. The representative mass spectra of the PDCI group chosen for validation are provided in supplementary results (S2).

We performed one-way ANOVA for the statistical analysis using Perseus software, with p < 0.05 as significant. The data pre-filter was based on presence of valid values in at least 3 samples in each group. A heat map (fig. 3A) depicts the hierarchical clustering of differentially expressed proteins representing LFQ intensity values, which are equivalent to the expression of proteins. Venn diagrams were plotted using bioinformatics and evolutionary genomics to cluster the overlapping and/or uniquely altered proteins (fig. 3B, 3C). The distribution of LFQ differences (log fold change) (x-axis) and p-value (y-axis) are represented as volcano plots [fig. 4(A-C)] on Perseus following ANOVA (FDR<0.05). A threshold of >1.5 fold change (2*^log fold change^) was set as significant upregulation and that of <0.5 was considered as significant down-regulation. Differentially expressed proteins were subjected to gene ontology and gene enrichment analysis using FunRich version 3.1.3. [20]

**Fig. 1:**
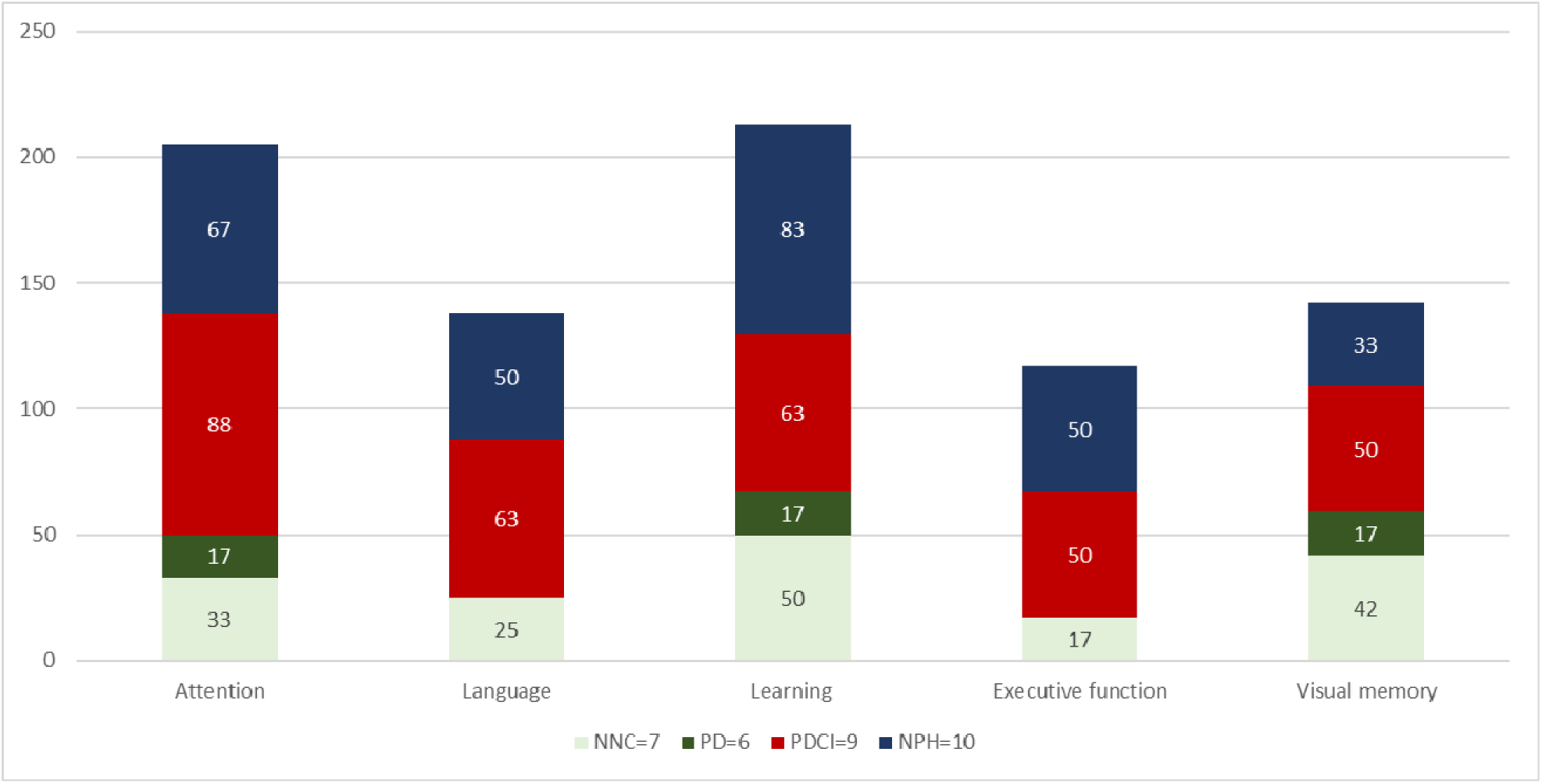
Histogram representing percentage of patients showing impairment in cognitive domains viz. attention, language, learning, executive and visuospatial domains. Number of subjects/patients enrolled NNC = 11; PDCI = 10; PD = 6, NPH = 10. Note that 88 % PDCI patients showed impairment in attention; 63% in language and learning; 50% in both executive and visuospatial function. A higher number (83%) of NPH patients showed impairment in learning. Among the PD patients 17 % showed impairment in attention, learning & visuospatial domains. NNC also showed higher (42%) impairment in visuospatial domain.

**Fig. 2:**
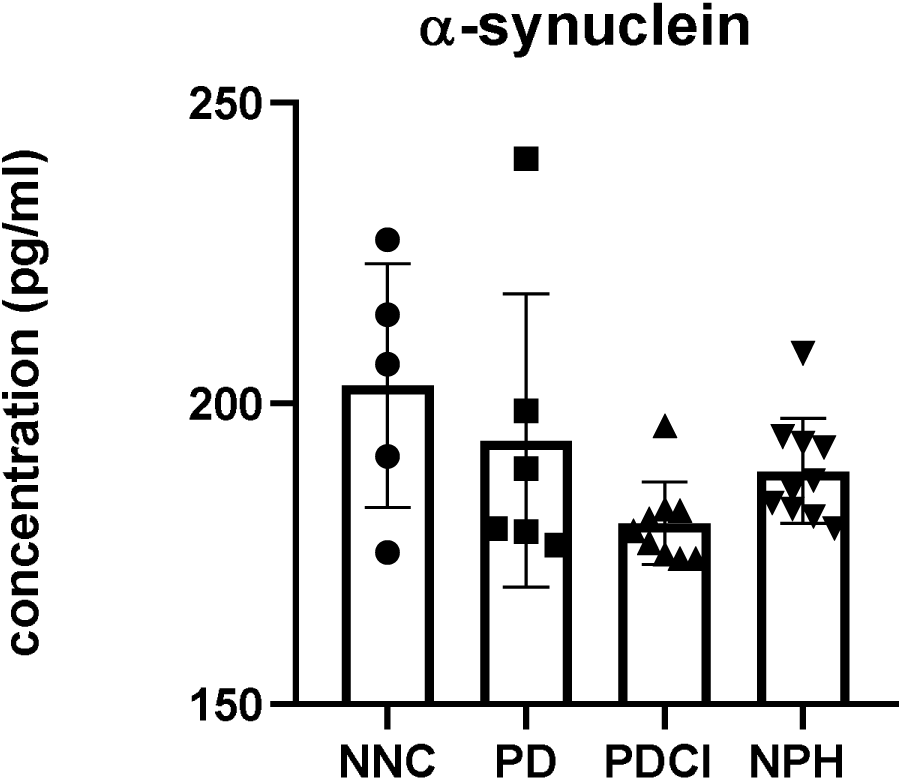
Histogram depicting individual values for ELISA-based estimation of α-synuclein in CSF of NNC (n=6), PD (n=6), PDCI (n=9) & NPH (n=10). Note mild, non-significant, decrease in its levels in PDCI-CSF [PDCI vs. NNC p=0.0762; PDCI vs. PD; p=0.8735; PDCI vs. NPH p=0.1735].

**Fig 3:**
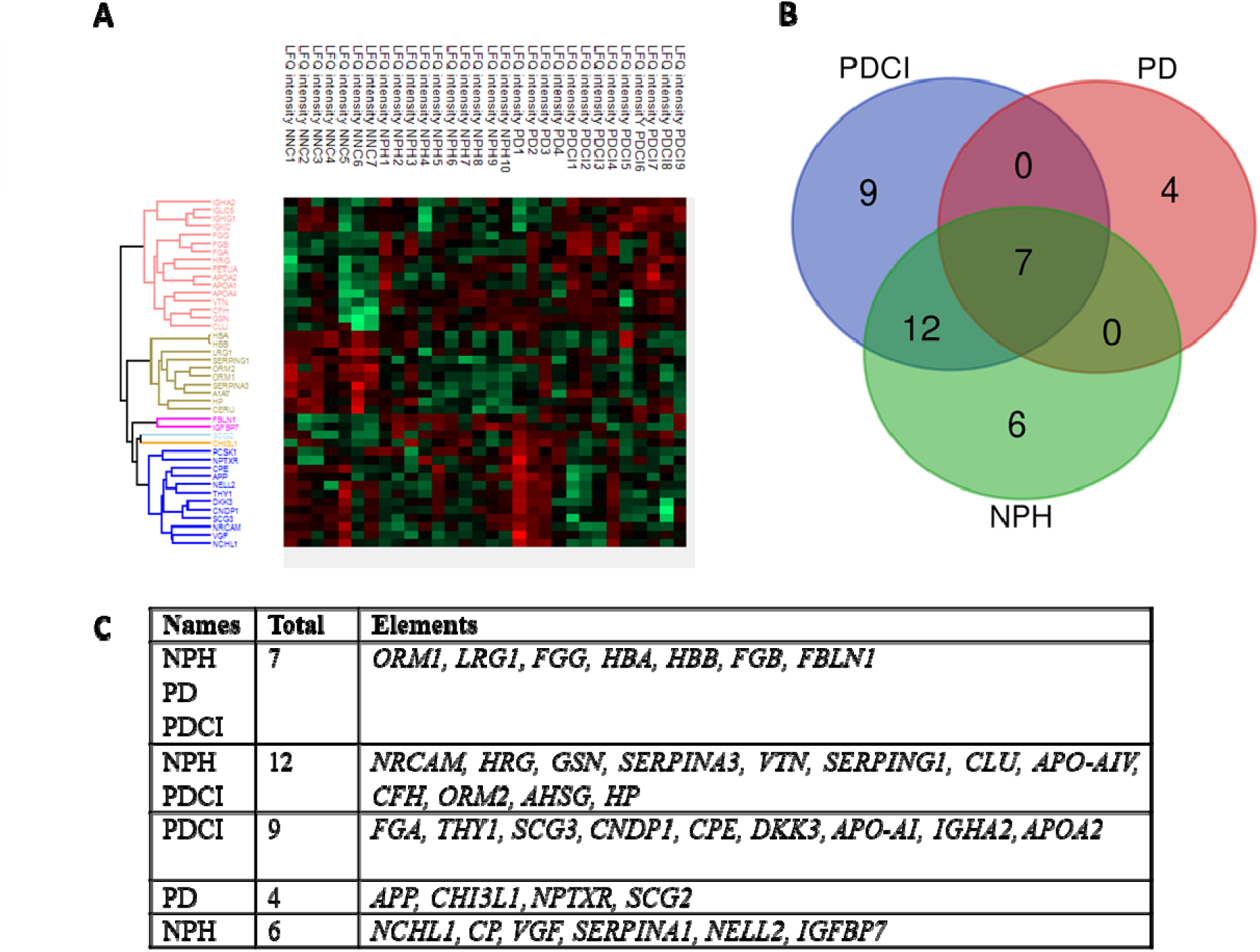
Heat map showing hierarchical expression of LFQ intensities depicting altered protein expression in PDCI, NPH & PD. (A) Of the 282 identified proteins 42 proteins were differentially altered. (B) Venn diagram showing 9, 4 and 6 proteins being uniquely altered in PDCI, PD and NPH respectively, whereas 7 proteins overlapped in PDCI and NPH. (C) Table showing the list of unique and common proteins with altered expression in PD, PDCI and NPH.

**Fig 4:**
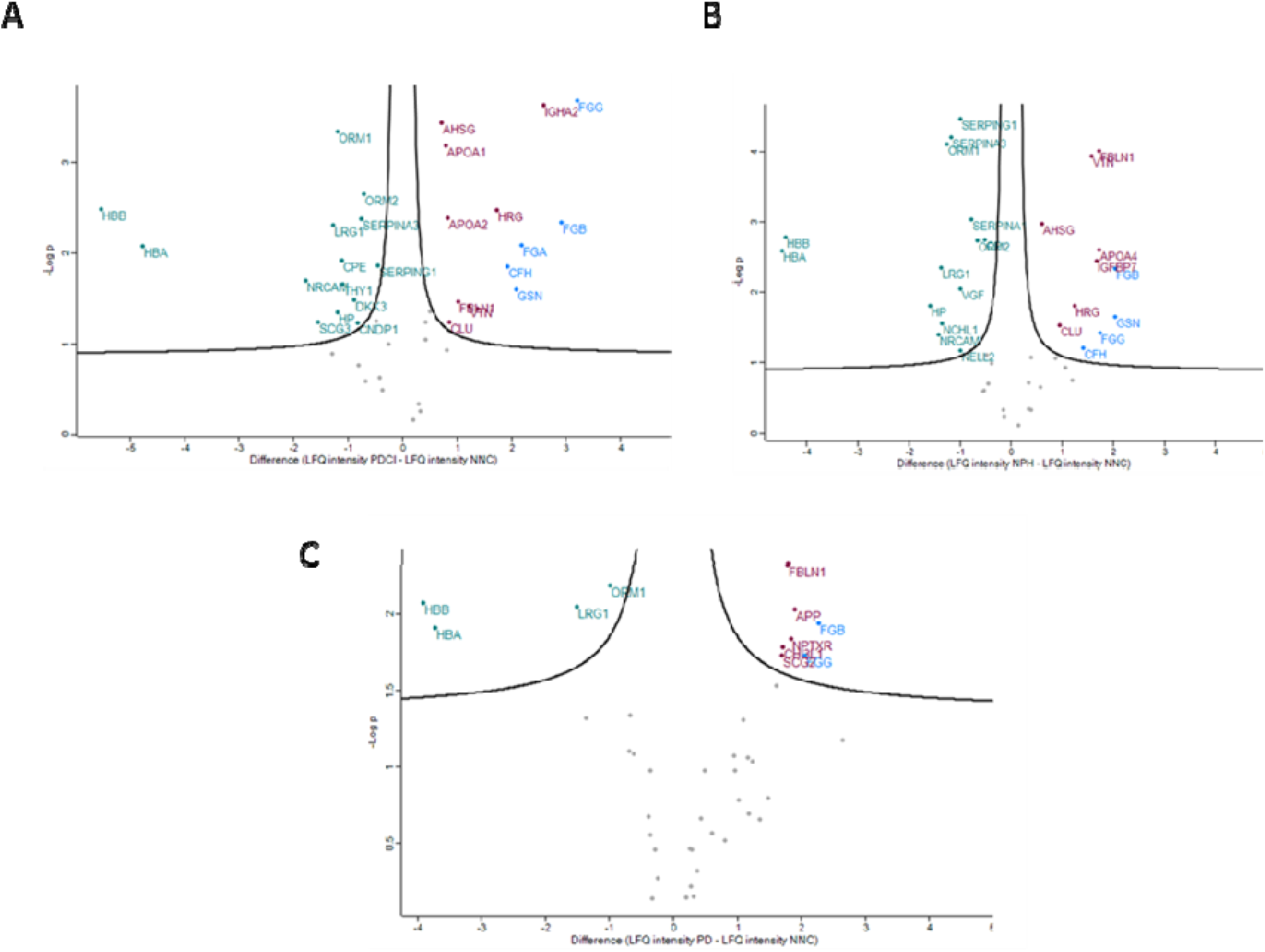
Volcano plots showing fold changes in LFQ intensities (A) PDCI vs NNC. (B) NPH vs NNC (C) PD vs NNC. Ordinate represents log p-values, and the abscissa represents the log fold changes. Significantly down-regulated proteins are highlighted in green (left quadrant) whereas significantly upregulated proteins highlighted in red (right quadrant). Proteins highlighted in blue were chosen for validation by ELISA. **Note:** Names of genes (italicized) and proteins have been used interchangeably.

### ELISA

We estimated the levels of α-synuclein in the CSF of patients using ELISA technique, since it is a reported biomarker for PD and PDCI (NNC, n=5-6; PD, n=6; PDCI, n= 8-11; NHP, n=7-10). Further, we also validated the proteomics findings by ELISA, for 5 significantly upregulated proteins viz. human fibrinogen, gelsolin, CFAH, ApoA-IV and ApoA-I in the CSF. The concentrations were measured in each of the CSF samples using commercially available ELISA kits [α-synuclein: ab210973 Abcam UK; (Human Fibrinogen: ab208036 Abcam UK, Human Gelsolin: ITRP09965 Immunotag, USA, Human Apolipoprotein AIV: ab214567 Abcam UK, Human Apolipoprotein AI: ab108804 Abcam UK, Human Complement factor H: ab252359 Abcam UK)].

### Statistical analysis

The ELISA data was analyzed using non-parametric Kruskal Wallis test followed by Dunn’s test for post hoc analysis, by Graphpad Prism (version 8). We further correlated the ELISA test results with the neuropsychological test percentiles by group-wise Kendall’s correlation and LFQ intensities by Spearman’s correlation using SPSS software.

## Results

### Demographic and clinical characteristics

Demographic analysis (Table. 1) showed higher number of male subjects/patients when compared to females. The mean disease duration for PDCI was 3 years post PD onset. The MOCA scores of PDCI patients confirmed cognitive deficit as they fell below the cut-off score of 26 within a range of (8-24). The median OFF state and ON state scores of PDCI were 47.25 and 41.5, respectively.

### Neuropsychological and radiological assessment findings

The battery of neuropsychological tests revealed that 88% PDCI patients had impairment in attention domain, 63% in language & learning domains, 50% in executive and 50% in visual memory domain (Table. 1). Interestingly, in comparison to PDCI a higher percentage (83%) of NPH patients showed impairment in learning whereas equal numbers showed impaired executive function. In PD, only around 17% patients showed cognitive impairment in attention, learning and visual memory domains, although in NNCs it was approximately 50% in learning and 42% visual memory tasks. Radiological findings by MRI supported the diagnosis of NPH due to the presence of ventricular dilatation, acute callosal angle between the lateral ventricles on coronal MRI images and in cases of PD by the fading of the swallow tail sign suggestive of nigrosome 1 alteration [21].

### Identification of proteins by unbiased Mass Spectrometry

The identified proteins (282) are listed in supplementary data (S3-excel). Amongst the 42 differentially expressed proteins, the highly upregulated ones in PDCI were fibrinogen α, β and γ chains; gelsolin, *CFH*, ApoA-IV and ApoA-I. Besides, a few proteins showed disease specific alterations (Table 6). The proteins uniquely altered in PDCI (with fold changes in parenthesis) were *IGHA2* (6.00), *FGA* (4.53), *APO-AI* (1.77), *APO-AII* (1.73), *CNDP1* (0.56), *DKK3* (0.54), *THY-1* (0.46), *CPE* (0.46), *SCG3* (0.34). Those specific for PD were *(APP)* (3.74), *NPTXR* (3.59), *CHI3L1* (3.27), *SCG2* (3.24). NPH also had specifically altered proteins e.g. *IGFBP7* (3.19), *CP* (0.70), *ORM2* (0.58), VGF nerve growth factor (0.50), *NRCAM* (0.39).

**Table 3:** Altered proteins in PDCI vs NNC. Note the LFQ difference and the fold change of proteins (Log fold = LFQ difference).

| SL No. | Gene names | Protein names | LFQ difference | Fold change | -log (p value) |
| --- | --- | --- | --- | --- | --- |
| 1 | *FGG* | Fibrinogen $\beta$ chain | 3.206306 | 9.229844 | 3.679611 |
| 2 | *FGB* | Fibrinogen $\gamma$ chain | 2.906603 | 7.498507 | 2.340156 |
| 3 | *IGHA2* | Immunoglobulin heavy constant α2 | 2.588646 | 6.015339 | 3.627602 |
| 4 | *FGA* | Fibrinogen $\alpha$ chain | 2.180321 | 4.532543 | 2.089083 |
| 5 | *GSN* | Gelsolin | 2.088767 | 4.253843 | 1.596505 |
| 6 | *CFH* | Complement factor H | 1.920743 | 3.786179 | 1.855669 |
| 7 | *HRG* | Histidine-rich glycoprotein | 1.729997 | 3.317271 | 2.477776 |
| 8 | *APOA-IV* | Apolipoprotein A4 | 1.374554 | 2.592878 | 1.387519 |
| 9 | *VTN* | Vitronectin | 1.217881 | 2.326048 | 1.420125 |
| 10 | *FBLN1* | Fibulin 1 | 1.021255 | 2.029684 | 1.460066 |
| 11 | *CLU* | Clusterin | 0.861097 | 1.816419 | 1.238028 |
| 12 | *APOA-II* | Apolipoprotein A2 | 0.829822 | 1.777466 | 2.390923 |
| 13 | *APOA-I* | Apolipoprotein A1 | 0.792492 | 1.732064 | 3.18239 |
| 14 | *AHSG* | Fetuin-A | 0.715055 | 1.641546 | 3.441942 |
| 15 | *SERPING1* | C1- inhibitor | -0.45606 | 0.728973 | 1.86647 |
| 16 | *ORM2* | Alpha-1-acid glycoprotein 2 precursor | -0.71635 | 0.608635 | 2.646635 |
| 17 | *SERPINA3* | Alpha-1-antichymotrypsin | -0.76075 | 0.590189 | 2.381479 |
| 18 | *CNDP1* | Carnosine Dipeptidase 1 | -0.81652 | 0.567808 | 1.231069 |
| 19 | *DKK3* | Dickkopf 3 | -0.88738 | 0.540594 | 1.486255 |
| 20 | *THY1* | Thy-1 membrane glycoprotein | -1.09575 | 0.467892 | 1.661783 |
| 21 | *CPE* | Carboxypeptidase E (CPE)<br>Or Neurotrophic factor-α1 | -1.1076 | 0.464066 | 1.922986 |
| 22 | *Orosomucoid 1 or ORM1* | Alpha-1-acid glycoprotein 1 precursor | -1.19579 | 0.436549 | 3.336183 |
| 23 | *HP* | Haptoglobin | -1.1973 | 0.436089 | 1.34499 |
| 24 | *LRG1* | Leucine-rich alpha-2-glycoprotein | -1.27958 | 0.411914 | 2.305596 |
| 25 | *SCG3* | Secretogranin III | -1.5509 | 0.341297 | 1.244947 |
| 26 | *NRCAM* | Neuronal cell adhesion molecule | -1.78338 | 0.290502 | 1.699357 |
| 27 | *HBA* | Haemoglobin $\alpha$ chain | -4.77007 | 0.036649 | 2.074899 |
| 28 | *HBB* | Haemoglobin $\beta$ chain | -5.52946 | 0.02165 | 2.482579 |

**Table 4:** Altered expression proteins in PD vs NNC. Note the LFQ difference and the fold change of proteins.

| SL No. | Gene names | Protein names | LFQ difference | Fold change | -log (p value) |
| --- | --- | --- | --- | --- | --- |
| 1 | *FGB* | Fibrinogen $\beta$ chain | 2.268839257 | 4.819352 | 1.942482 |
| 2 | *FGG* | Fibrinogen $\gamma$ chain | 2.054453577 | 4.153863 | 1.729726 |
| 3 | *APP* | Amyloid precursor protein | 1.90311483 | 3.740198 | 2.034483 |
| 4 | *NPTXR* | Neuronal pentraxin receptor | 1.845703568 | 3.594282 | 1.840862 |
| 5 | *FBLN1* | Fibulin 1 | 1.775449889 | 3.423448 | 2.321667 |
| 6 | *CHI3L1* | Chitinase-3-like protein 1 | 1.709992647 | 3.271592 | 1.78289 |
| 7 | *SCG2* | Secretogranin-2 precursor | 1.697182076 | 3.24267 | 1.734019 |
| 8 | *HRG* | Histidine-rich glycoprotein | 1.60633564 | 3.044775 | 1.534191 |
| 9 | *ORM1* | Alpha-1-acid glycoprotein 1 precursor | -0.995535033 | 0.50155 | 2.191851 |
| 10 | *LRG1* | Leucine-rich alpha-2-glycoprotein | -1.507603986 | 0.351695 | 2.052224 |
| 11 | *HBA* | Haemoglobin $\alpha$ chain | -3.721756969 | 0.075795 | 1.911425 |
| 12 | *HBB* | Haemoglobin $\beta$ chain | -3.911985602 | 0.066432 | 2.072433 |

**Table 5:** Altered expression of proteins in NPH vs NNC. Note the LFQ difference and the fold change of proteins.

| SL No. | Gene names | Protein names | LFQ difference | Fold change | -log (p value) |
| --- | --- | --- | --- | --- | --- |
| 1 | *FGB* | Fibrinogen $\beta$ chain | 2.034424 | 4.096592 | 2.337591 |
| 2 | *GSN* | Gelsolin | 2.032031 | 4.089801 | 1.6519 |
| 3 | *FGG* | Fibrinogen $\gamma$ chain | 1.745789 | 3.353781 | 1.425847 |
| 4 | *FBLN1* | Fibulin 1 | 1.73209 | 3.322087 | 4.017884 |
| 5 | *APOA-IV* | Apolipoprotein A-IV | 1.728503 | 3.313838 | 2.611286 |
| 6 | *IGFBP7* | Insulin-like growth factor-binding protein | 1.675269 | 3.193788 | 2.447808 |
| 7 | *VTN* | Vitronectin | 1.578295 | 2.986166 | 3.937421 |
| 8 | *CFH* | Complement Factor H | 1.407013 | 2.651875 | 1.211394 |
| 9 | *HRG* | Histidine-rich glycoprotein | 1.239616 | 2.361356 | 1.812513 |
| 10 | *CLU* | Clusterin | 0.958206 | 1.942892 | 1.538422 |
| 11 | *AHSG* | Fetuin A | 0.59755 | 1.513144 | 2.973582 |
| 12 | *CP* | Ceruloplasmin | -0.50918 | 0.70262 | 2.754692 |
| 13 | *ORM2* | Alpha-1-acid glycoprotein 2 precursor | -0.66813 | 0.629323 | 2.739543 |
| 14 | *SERPINA1* | Alpha-1 antitrypsin | -0.78105 | 0.581944 | 3.041422 |
| 15 | *SERPING1* | C1- inhibitor | -0.98824 | 0.504093 | 4.470284 |
| 16 | *NELL2* | Neural EGFL like 2 | -0.99017 | 0.503417 | 1.170732 |
| 17 | *VGF* | VGF nerve growth factor | -0.992 | 0.50278 | 2.054869 |
| 18 | *SERPINA3* | Alpha-1-antichymotrypsin | -1.16581 | 0.445714 | 4.211002 |
| 19 | *ORM1* | Alpha-1-acid glycoprotein 1 precursor | -1.25082 | 0.420211 | 4.105446 |
| 20 | *NCHL1* | Neural cell adhesion molecule L1 like protein | -1.35802 | 0.390118 | 1.564199 |
| 21 | *LRG1* | Leucine-rich alpha-2-glycoprotein | -1.36901 | 0.387157 | 2.349068 |
| 22 | *NRCAM* | Neuronal cell adhesion molecule | -1.41429 | 0.375195 | 1.399183 |
| 23 | *HP* | Haptoglobin | -1.58972 | 0.332236 | 1.815579 |
| 24 | *HBB* | Haemoglobin $\alpha$ chain | -4.41316 | 0.046936 | 2.780855 |
| 25 | *HBA* | Haemoglobin $\beta$ chain | -4.48538 | 0.044644 | 2.590615 |

**Table 6:** Proteins that were uniquely altered in PD, PDCI & NPH along with their respective fold changes

| SL No. | PD | PDCI | NPH |
| --- | --- | --- | --- |
| 1 | Amyloid precursor protein<br>3.74 | Immunoglobulin heavy constant $\alpha 2$<br>6.00 | Insulin-like growth factor-binding protein 7<br>3.19 |
| 2 | Neuronal pentraxin receptor<br>3.59 | Fibrinogen $\alpha$ chain<br>4.53 | Ceruloplasmin<br>0.70 |
| 3 | Chitinase-3-like protein 1<br>3.27 | Apolipoprotein A2<br>1.77 | Alpha-1 antitrypsin<br>0.58 |
| 4 | Secretogranin-2 precursor<br>3.24 | Apolipoprotein A1<br>1.73 | VGF Nerve growth factor<br>0.50 |
| 5 |  | Carnosine Dipeptidase 1<br>0.56 | Neural cell adhesion molecule L1 like protein<br>0.39 |
| 6 |  | Dickkopf 3<br>0.54 |  |
| 7 |  | Thy-1 membrane glycoprotein<br>0.46 |  |
| 8 |  | Carboxypeptidase E (CPE)<br>0.46 |  |
| 9 |  | Secretogranin 3 precursor<br>0.34 |  |

### Upregulated proteins that may predict cognitive impairment in PD

ELISA findings suggested that α-synuclein levels were mildly lower in PDCI when compared to NNC, PD and NPH (fig. 2; NNC vs PDCI p = 0.0762, PD vs PDCI p = 0.8735 and NPH vs PDCI p = 0.1735). We further validated the MS/MS findings by targeted ELISA for each protein. A mild increase in the fibrinogen levels in PDCI was noted in comparison to NNC (p = 0.1575); whereas the NPH samples showed significantly higher levels (fig. 5A; **p = 0.0032 NNC vs NPH). Gelsolin was significantly higher in PD (fig. 5B; *p = 0.0109 NNC vs PD) compared to NNC. Interestingly, its levels were low in PDCI (fig. 5B; *p = 0.0368 PD vs PDCI; ANOVA), but failed to show significance following post-hoc analysis (Dunn’s correction).

**Fig. 5:**
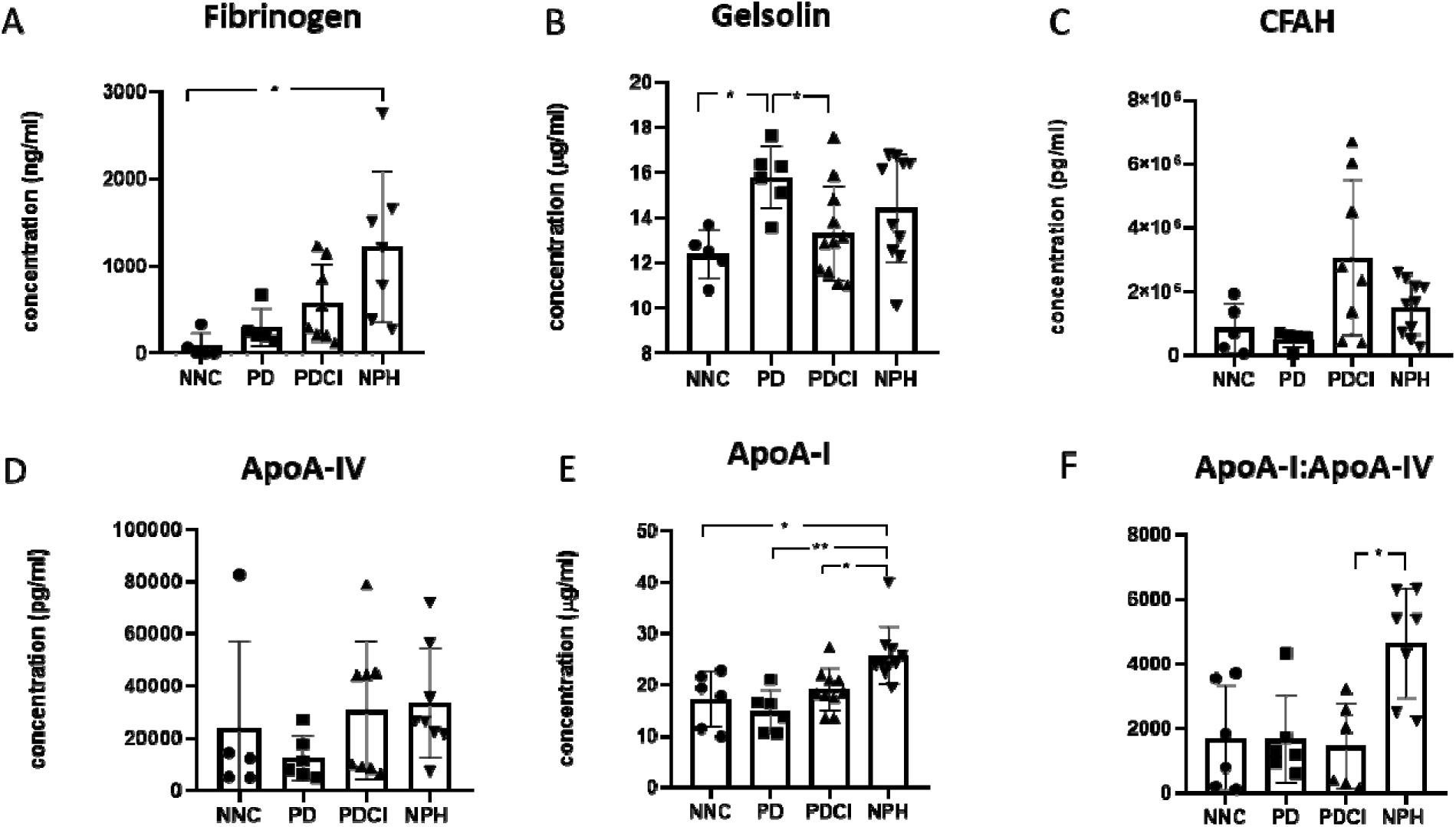
Histogram with individual values representing ELISA-derived protein concentrations in patient CSF. A. Note the significantly high fibrinogen levels in NPH (NNC vs NPH **p = 0.0032). B. Note the significantly higher level of gelsolin in PD (NNC vs PD *p = 0.0109) and lower level in PDCI (PD vs PDCI *p = 0.0368). C & D: No significant difference was observed in CFAH and ApoA-IV respectively. Note the mildly high CFAH in PDCI (PDCI vs. PD, p = 0.1030) E. ApoA-I is significantly more in NNC vs NPH *p= 0.0287, PD vs NPH **p = 0.0012, PDCI vs NPH *p = 0.0395. F. The ratio of ApoA-I: ApoA-IV was significantly higher in NPH (PDCI vs NPH *p= 0.0456).

CFAH showed a mild, non-significant, increase in PDCI compared to PD p=0.1030 (fig. 5C). We found a mild, non-significant, upregulation of ApoA-IV in PDCI and NPH when compared to PD (fig. 5D). The levels of ApoA-I were significantly lower in PDCI than NPH (fig.5E; *p=0.0395NPH vs PDCI). Interestingly, a significantly high ratio of ApoA-I:ApoA-IV proteins was noted in PDCI in comparison to PD and NPH (fig. 5F; *p=0.0456 NPH vs PDCI).

### Gene ontology-based analysis

The proteins altered in PDCI-CSF were associated with various biological processes as well as molecular and cellular components [fig. 6a (A-C); (table. 7 (A-C)]. In PDCI, 32% of the altered proteins (*FGG, FGB, FGA, SERPING1, SERPINA3, CNDP1, CPE, SCG3*) were associated with protein metabolism, (12%) (*AHSG, ORM2, ORM1*) with immune activity and 64% (*FGG, FGB, FGA, GSN, CFH, HRG, VTN, CLU, APOA-II, APOA-I, AHSG, SERPING1, SERPINA3, THY1, HP, HBB*) showed exosome(s) association and 84 % (*FGG, FGB, FGA, GSN, CFH, HRG, VTN, FBLN1, CLU, APOA-II, APOA-I, AHSG, SERPING1, ORM2, SERPINA3, DKK3, ORM1, HP, LRG1, SCG3, HBB*) were extracellular. In PD, [fig. 6b (A-C), table.9 (A-C)] 27.3 % of the altered proteins belonged to protein metabolism (*FGB, FGG, SCG2*), 9% were associated with apoptosis (*HRG*), 18.2% belonged to the extracellular matrix structure (*FBLN1, CHI3L1*); and 18.2 % belonged to cytoskeleton (*FGB, FGG*) as well as fibrinogen complex (*FGB, FGG*). In NPH [fig 6c.(A-C); table.9(A-C)] 22.7% altered proteins were associated with protein metabolism (*FGB, FGG, SERPINA1, SERPING1, SERPINA3*) and immune response (*CFH, CLU, ORM2, ORM1, HP*); 13.6 % were protease inhibitors (*IGFBP7, NRCAM*), defense/ immune protein activity (*AHSG, ORM2, ORM1*) and 18.2% were cytoskeletal proteins.

**Fig. 6:**
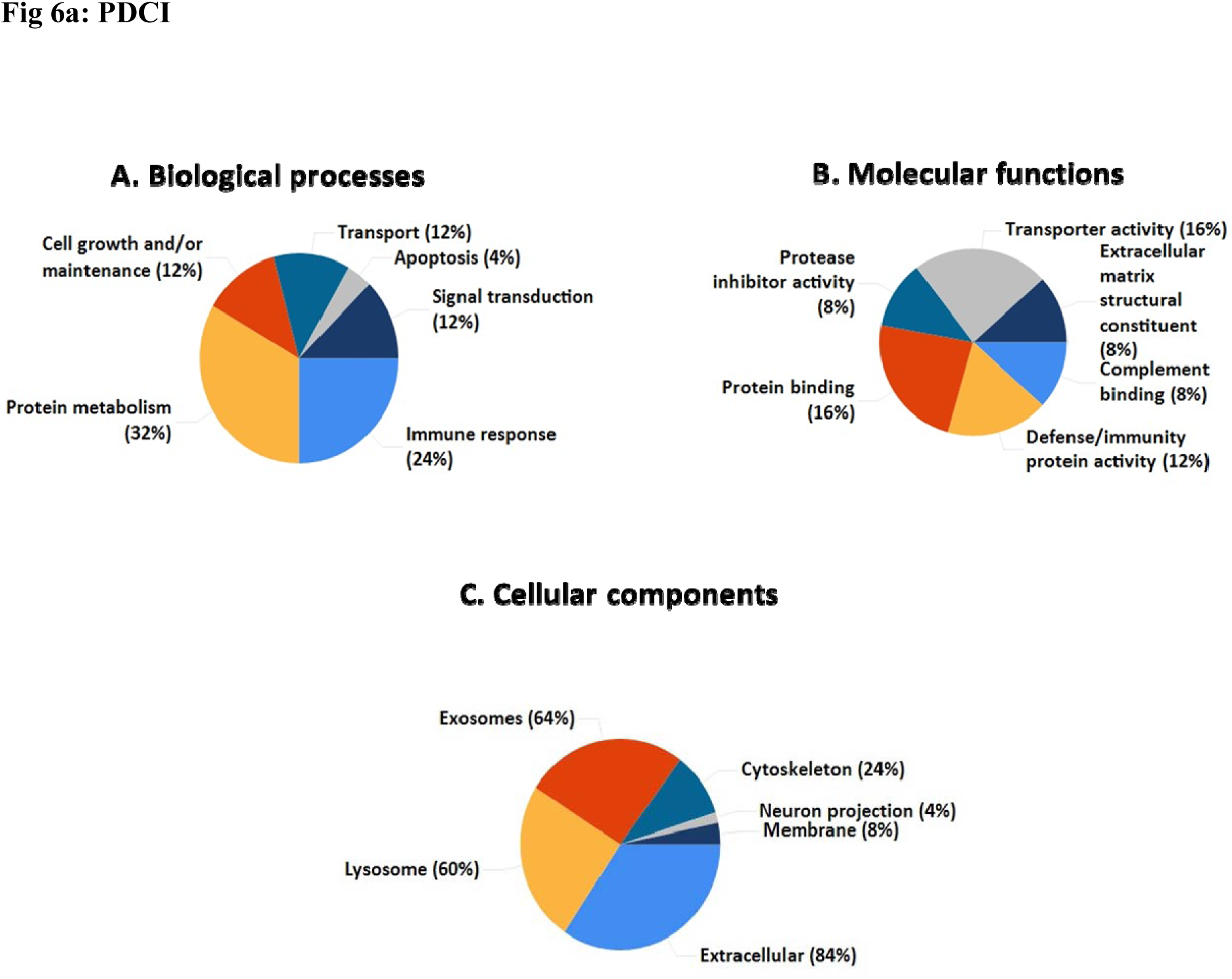
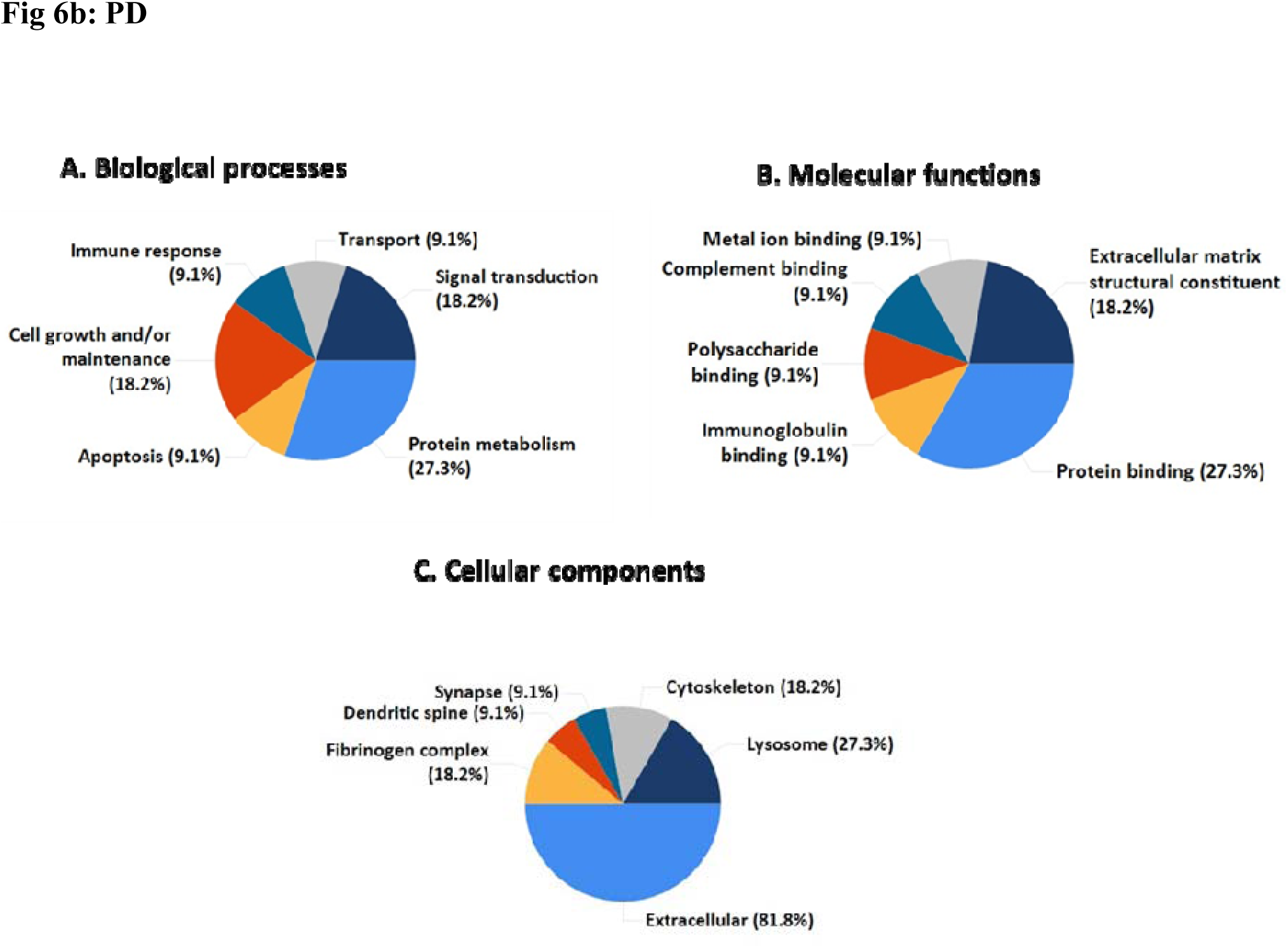
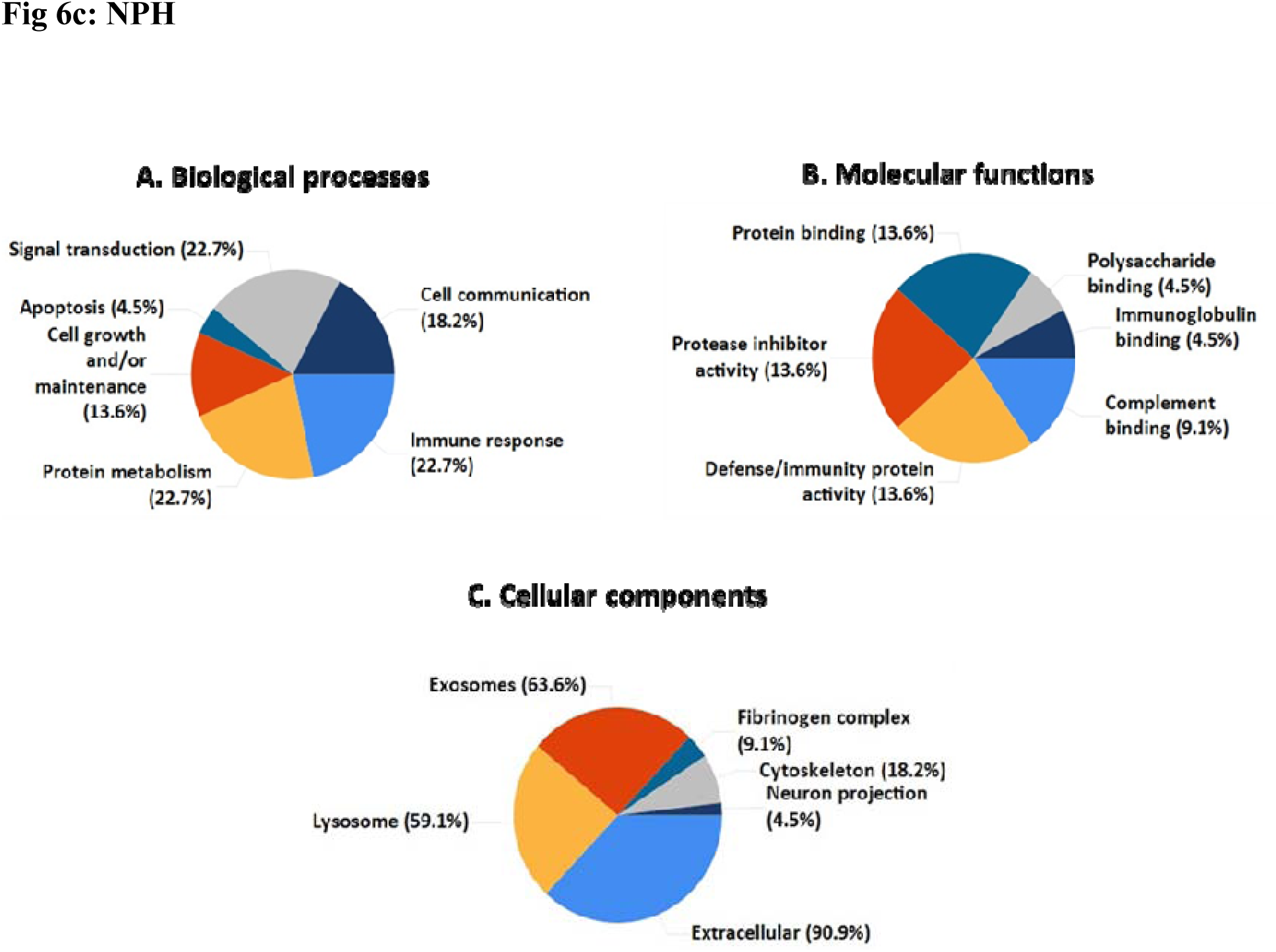
Pie charts representing the gene ontology-based percentage of altered proteins identified by the FunRich software in the different study groups. **Fig. 6a:** PDCI A. Biological processes B. Molecular functions C. Cellular components. 6 entities were chosen for each category in the pie chart display. Out of 28 gene products 25 were mapped. **Fig. 6b: PD** A. Biological processes B. Molecular functions C. Cellular components. 7 biological processes are represented in A. whereas B. & C. present 6 molecular functions and cellular components. **Fig. 6c:** NPH A. Biological processes B. Molecular functions C. Cellular components. 6 entities were chosen for each category in the pie chart display.

**Table 7:** List of processes affected and corresponding gene products in PDCI as per gene ontology A. biological process, B. molecular function and C. cellular component. 6 aspects per category have been represented.

|  |  |
| --- | --- |
| **A. Biological processes** | **Genes mapped** |
| Protein metabolism | *FGG, FGB, FGA, SERPING1, SERPINA3, CNDP1, CPE, SCG3* |
| Cell growth and/or maintenance | *GSN, VTN, FBLN1* |
| Transport | *APOA2, APOA1, HBB* |
| Signal transduction | *AHSG, DKK3, NRCAM* |
| Immune response | *CFH, CLU, ORM2, THY1, ORM1, HP* |
| Apoptosis | *HRG* |
| **B. Molecular functions** | **Genes mapped** |
| Protein binding | *FGG, FGB, FGA, HRG* |
| Defense/Immunity | *AHSG, ORM2, ORM1* |
| Protease inhibitor | *SERPING1, SERPINA3* |
| Complement binding | *CLU* |
| Transporter activity | *APOA1, APOA2, HP, HBB* |
| Immunoglobulin binding | *HRG* |
| **C. Cellular components** | **Genes mapped** |
| Exosomes | *FGG, FGB, FGA, GSN, CFH, HRG, VTN, CLU, APOA2, APOA1, AHSG, SERPING1, SERPINA3, THY1, HP, HBB* |
| Lysosome | *FGA, GSN, CFH, VTN, CLU, APOA2, APOA1, AHSG, SERPING1, SERPINA3, THY1, ORM1, HP, NRCAM, HBB* |
| Extracellular | *FGG, FGB, FGA, GSN, CFH, HRG, VTN, FBLN1, CLU, APOA2, APOA1, AHSG, SERPING1, ORM2, SERPINA3, DKK3, ORM1, HP, LRG1, SCG3, HBB* |
| Membrane | *THY1, LRG1* |
| Neuron projection | *NRCAM* |

**Table 8:**
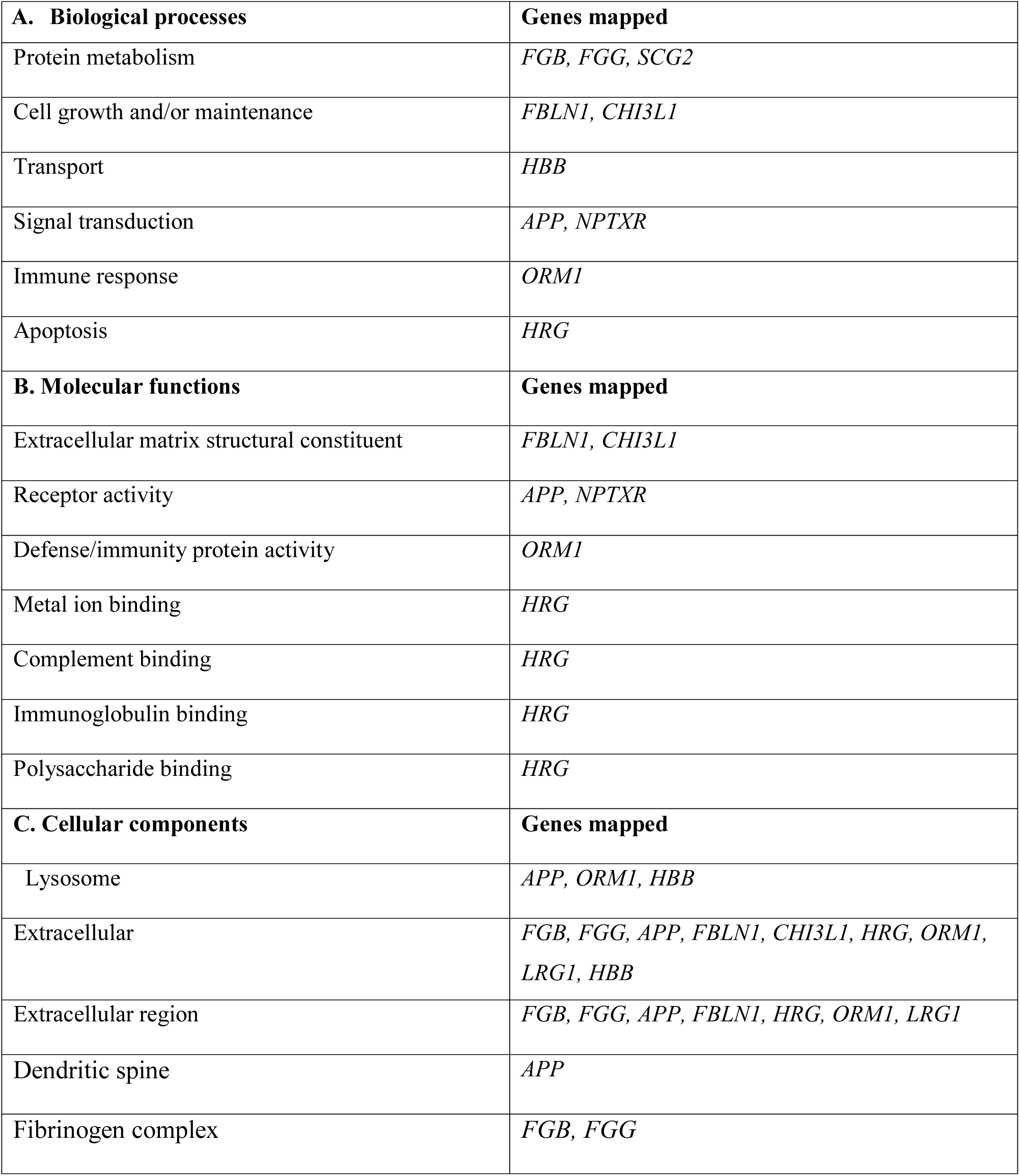
List of processes affected and corresponding gene products in PD as per gene ontology. A. biological process, B. molecular function and C. cellular component. 6 aspects per category have been represented.

| **A. Biological processes** | **Genes mapped** |
| --- | --- |
| Protein metabolism | *FGB, FGG, SCG2* |
| Cell growth and/or maintenance | *FBLN1, CHI3L1* |
| Transport | *HBB* |
| Signal transduction | *APP, NPTXR* |
| Immune response | *ORM1* |
| Apoptosis | *HRG* |
| **B. Molecular functions** | **Genes mapped** |
| Extracellular matrix structural constituent | *FBLN1, CHI3L1* |
| Receptor activity | *APP, NPTXR* |
| Defense/immunity protein activity | *ORM1* |
| Metal ion binding | *HRG* |
| Complement binding | *HRG* |
| Immunoglobulin binding | *HRG* |
| Polysaccharide binding | *HRG* |
| **C. Cellular components** | **Genes mapped** |
| Lysosome | *APP, ORM1, HBB* |
| Extracellular | *FGB, FGG, APP, FBLN1, CHI3L1, HRG, ORM1, LRG1, HBB* |
| Extracellular region | *FGB, FGG, APP, FBLN1, HRG, ORM1, LRG1* |
| Dendritic spine | *APP* |
| Fibrinogen complex | *FGB, FGG* |

**Table 9:** List of processes affected and corresponding gene products in NPH as per gene ontology A. biological process, B. molecular function and C. cellular component. 6 aspects per ontology A. biological process, B. molecular function and C. cellular component. 6 aspects per

| **A. Biological process** | **Genes mapped** |
| --- | --- |
| Protein metabolism | *FGB, FGG, SERPINA1, SERPING1, SERPINA3* |
| Cell growth and/or maintenance | *GSN, FBLN1, VTN* |
| Cell communication | *IGFBP7, AHSG, NELL2, VGF* |
| Signal transduction | *IGFBP7, AHSG, NELL2, VGF, NRCAM* |
| Immune response | *CFH, CLU, ORM2, ORM1, HP* |
| Apoptosis | *HRG* |
| **B. Molecular function** | **Genes mapped** |
| Protease inhibitor activity | *SERPINA1, SERPING1, SERPINA3* |
| Defense/immunity protein activity | *AHSG, ORM2, ORM1* |
| Protein binding | *FGB, FGG, HRG* |
| Complement binding | *CFH, HRG* |
| Immunoglobulin binding | *HRG* |
| Polysaccharide binding | *HRG* |
| **C. Cellular component** | **Genes mapped** |
| Exosomes | *FGB, GSN, FGG, VTN, CFH, HRG, CLU, AHSG, CP, SERPINA1, SERPING1, SERPINA3, HP, HBB* |
| Lysosome | *GSN, VTN, CFH, CLU, AHSG, CP, SERPINA1, SERPING1, SERPINA3, ORM1, NRCAM, HP, HBB* |
| Extracellular | *FGB, GSN, FGG, FBLN1, IGFBP7, VTN, CFH, HRG, CLU, AHSG, CP, ORM2, SERPINA1, SERPING1, VGF, SERPINA3, ORM1, LRG1, HP, HBB* |
| Cytoskeleton | *FGB, GSN, FGG, CLU* |
| Fibrinogen complex | *FGB, FGG* |
| Neuron projection | *NRCAM* |

### Statistical correlations with ELISA

Spearman’s correlation showed a positive correlation between the LFQ intensity of *FGA* and the ELISA-derived concentration of native fibrinogen (ρ = 0.750, *p = 0.020). Similarly, CFAH correlated positively with native fibrinogen (ρ = 0.460, *p = 0.036) (Table.10). We further assessed whether the protein levels correlated with the neuropsychological outcome. A group wise Kendall’s correlation analysis (Table. 11) showed a positive correlation between ELISA-derived CFAH concentration and digit vigilance test percentile (r = 0.949, *p value = 0.023). On a similar note the ratio of ApoA-I:ApoA-IV showed a positive correlation (r = 0.894, *p = 0.037) while ApoA-IV showed a negative correlation with ANT percentile (r = −0.894, *p = 0.037). ApoA-I correlated positively with verbal N-BACK test (r = 0.683, *p = 0.033) while fibrinogen showed a negative correlation with WCST (r = −0.788, *p = 0.032).

**Table 10.** LFQ intensities vs ELISA protein concentration. Spearman’s rho correlation between LFQ intensities and ELISA based concentrations of the highly up-regulated proteins in PDCI. Note a positive correlation (*) between LFQ intensity of highly up-regulated proteins in PDCI. Note a positive correlation (*) between LFQ intensity of showed positive correlation with the concentration of fibrinogen native protein.

|  |  | ELISA |  |  |  |  |  |
| --- | --- | --- | --- | --- | --- | --- | --- |
|  |  | Fibrinogen<br>(ng/ml) | Gelsolin<br>(ug/ml) | CFAH<br>(pg/ml) | ApoA-I<br>(ug/ml) | ApoA-IV<br>(pg/ml) | ApoA-I:<br>ApoA-IV<sup>#</sup> |
| **LFQ intensities** |  |  |  |  |  |  |  |
| **FGA** | Correlation Coefficient | .750<sup>*</sup> | -.233 | -.150 | .200 | .143 | .024 |
|  | Sig. (2-tailed) | .020 | .546 | .700 | .606 | .736 | .955 |
|  | N | 9 | 9 | 9 | 9 | 8 | 8 |
| **FGB** | Correlation Coefficient | .420 | -.210 | .098 | -.280 | .539 | -.418 |
|  | Sig. (2-tailed) | .175 | .513 | .762 | .379 | .108 | .229 |
|  | N | 12 | 12 | 12 | 12 | 10 | 10 |
| **FGG** | Correlation Coefficient | .253 | -.253 | -.053 | -.226 | .363 | -.335 |
|  | Sig. (2-tailed) | .345 | .345 | .846 | .399 | .223 | .263 |
|  | N | 16 | 16 | 16 | 16 | 13 | 13 |
| **CFAH** | Correlation Coefficient | .460<sup>*</sup> | -.145 | .244 | -.004 | .408 | -.337 |
|  | Sig. (2-tailed) | .036 | .529 | .286 | .987 | .093 | .171 |
|  | N | 21 | 21 | 21 | 21 | 18 | 18 |
| **Gelsolin** | Correlation Coefficient | -.130 | .242 | -.229 | -.387 | .042 | -.065 |
|  | Sig. (2-tailed) | .575 | .291 | .319 | .083 | .868 | .798 |
|  | N | 21 | 21 | 21 | 21 | 18 | 18 |
| **APOA4** | Correlation Coefficient | .334 | -.213 | .257 | -.117 | .222 | -.253 |
|  | Sig. (2-tailed) | .139 | .354 | .260 | .614 | .376 | .311 |
|  | N | 21 | 21 | 21 | 21 | 18 | 18 |
| **APOA1** | Correlation Coefficient | .391 | -.273 | .319 | .021 | .245 | -.302 |
|  | Sig. (2-tailed) | .080 | .232 | .158 | .929 | .328 | .223 |
|  | N | 21 | 21 | 21 | 21 | 18 | 18 |

(<sup>#</sup> derived from pg/ml Apo-AI and Apo-AIV)

**Table 11:** Group wise Kendall’s correlation between neuropsychological test percentiles and ELISA derived concentration of proteins viz. Fibrinogen, CFAH, ApoA-I, ApoA-IV, ApoAI:ApoA-IV in PDCI. Significant correlation coefficients and p-values are denoted (*).

|  |  | ELISA |  |  |  |  |
| --- | --- | --- | --- | --- | --- | --- |
| Neuropsychological test | Comparisons | Fibrinogen (ng/ml) | CFAH (pg/ml) | ApoA-I (ug/ml) | ApoA-IV (pg/ml) | ApoAI: ApoA-IV |
| Digit vigilance total time (TT) | Correlation Coefficient | .316 | .949* | -.105 | .333 | -.333 |
|  | Sig. (2-tailed) | .448 | .023 | .801 | .602 | .602 |
|  | N | 5 | 5 | 5 | 3 | 3 |
| ANT | Correlation Coefficient | -.433 | .355 | .355 | -.894* | .894* |
|  | Sig. (2-tailed) | .154 | .244 | .244 | .037 | .037 |
|  | N | 8 | 8 | 8 | 5 | 5 |
| Verbal N back 2 error | Correlation Coefficient | 0.000 | .390 | .683* | -1.000* | 1.000* |
|  | Sig. (2-tailed) | 1.000 | .224 | .033 |  |  |
|  | N | 7 | 7 | 7 | 5 | 5 |
| WCST No. of categories completed | Correlation Coefficient | -.788* | -.215 | .358 | -.527 | .527 |
|  | Sig. (2-tailed) | .032 | .559 | .330 | .207 | .207 |
|  | N | 6 | 6 | 6 | 5 | 5 |

(\* derived from pg/ml Apo-AI and Apo-AIV)

Similar correlations were derived for NPH with (i) positive correlation between ApoA-I and MOCA scores (r = 0.506, *p = 0.046) (ii) a negative correlation between CFAH and COWA (CFAH vs COWA r = −0.567, *p = 0.049) and WCST percentile (r = −0.645, *p = 0.036); (iii) negative correlations of ApoA-IV with COWA (r = −0.619, *p = 0.036); ANT (r = −0.618, *p = 0.034), Verbal N-BACK 2 (r = −0.741, *p = 0.012) and AVLT immediate recall (IR) percentiles [(r = −0.732, *p = 0.029) (supplementary table. S1)].

## Discussion

This is the first study to suggest *FGA* as a possible biomarker to distinguish PDCI from NPH, based on LC-MS/MS analysis. Besides, as noted from ELISA-based observations, CFAH could potentially differentiate PDCI from PD. The correlations between the digit-vigilance and N-BACK tests with CFAH and ApoA-IV levels respectively, extend a diagnostic value to these proteins. No studies till date have reported an association of PDCI with these proteins. Further, attention deficit in PDCI and learning deficit in NPH patients [22] could be the non-invasive identifying features, while impairments in language and executive domains were common to both [22, 23]. Thus, a panel of proteins and neuropsychiatric assessments may assist prediction of cognitive decline in PDCI.

Various studies have reported a direct correlation between LB and cognitive impairment in PD [24, 25,]. Although α-synuclein is the main constituent of LBs, its reducing levels did not corroborate with UPDRS scores both cross-sectionally and longitudinally [26]. Our observation of mild decrease in α-synuclein thus matches earlier findings and led us on an exploratory study of non-targeted protein biomarker(s). An iTRAQ and multiple reactions monitoring study showed alterations in 16 proteins between PDD and PD [27]. Our observations on APO proteins in PDCI partly match theirs; albeit the variants are different. Some studies proposed alterations in clusterin, gelsolin, haptoglobin, and ApoA-I [28] in PD, while another showed serpin A1 to predict PDD at an early stage [29]. Our findings thus partly corroborate with these studies in relation to PDCI.

The specific upregulation of *FGA* in PDCI, vis-à-vis that of *FGB* and *FGG* in PD and NPH is intriguing. Fibrinogen is abundant in CSF in neurological disorders and traumatic CNS injury cases with blood brain barrier breach [30, 31]. It is synthesized by neurons and astroglia [32]. The absence of significant differences between controls and PDCI in ELISA findings may be due to the use of whole fibrinogen molecule. Yet, the positive correlation between LFQ intensities of *FGA* and whole fibrinogen, validate the use of latter. Marfà et al, considered lower levels of *FGA*-c-chain as a serum marker for hepatic fibrosis; while proposing that inflammation causes cleavage and release of fibrinogen fragments α, β and γ into the bloodstream or CSF. The cleavage sites depend on the disease etiology and mechanisms of protein degradation [33]. Gelsolin is present in the neurons, choroid plexus and CSF. Since it has anti-amyloidogenic properties [34], low levels in PDCI may represent a failed attempt at de-fibrillization whereas, the significant increase in PD may suggest a preserved function of severing of Aβ fibrils. The alternative pathway of the complement system that guides the innate immunity remains constitutively active to detect pathogens and altered self. CFAH is its prominent regulator [35]. In an ELISA-based study, complement protein C3 and regulatory protein factor H differentiated MSA from PD/AD. A correlation was noted between C3/Aβ and FH/Aβ and the severity of cognitive impairment in PD [36]. Elevated CFAH levels were noted in patients with MCI, without AD [37]. Similar elevation in our MS/MS readouts emphasizes its potential use in differentiating PD from PDCI. ELISA studies in a larger cohort may provide better insights.

Human ApoA-I, protein maintains cholesterol homeostasis in the CNS [38, 39]. Few in-vitro studies demonstrate a direct interaction between ApoA-I and APP in inhibiting Aβ plaque formation [40, 41]. Lower levels of ApoA-I in the plasma, was a potent risk factor for cognitive decline in MCI. A similar reduction in plasma/serum correlated well with severity of symptoms in AD [42, 43]. CSF collected at autopsy from AD cases too showed low levels [44]. The sizeable differences in its levels between our PDCI and NPH cases make it an interesting candidate. ApoA-IV is synthesized in limited amounts in the hypothalamus [45]. Although unaltered in an independent capacity it may still influence the pathogenesis as seen from the enhanced ratio of ApoA-I:ApoA-IV. Thus positive correlation between 1) CFAH and ApoA-I:ApoA-IV ratio with digit vigilance as well as ANT percentiles respectively 2) negative correlation between fibrinogen and verbal N-BACK along with WCST as well as 3) ApoA-IV levels and verbal N-BACK confirm their role in cognition-related pathology in PDCI.

Gene ontology studies in neurodegenerative disorders revealed that impaired protein degradation leads to abnormal aggregation of misfolded proteins like amyloid-β and α-synuclein, which may be subsequently transferred by exosomes. The proteins viz. *FGG, FGB, FGA, SERPING1, SERPINA3, CNDP1, CPE, SCG3 etc.* are associated with protein metabolism. The alterations in immune response proteins of the complement system and regulators like CFAH, clusterin and orosomucoid reflect compromised immune function. Exosomes being vesicles for intracellular transfer of cargos, the alterations in exosomal proteins (*FGG; FGB; FGA; GSN; CFH; HRG; VTN; CLU; APOA-II; APOA-I; AHSG; SERPING1; SERPINA3; THY1; HP; HBB*) affects protein logistics. Interestingly certain proteins modulate both protein metabolism as well as exosome related processes viz. (*FGG; FGB; FGA, SERPING1* S*SERPINA3*).

The exclusive downregulation of proteins like CNDP-1, DKK3, CPE and SCG3 precursor in PDCI, deserve detailed analysis. CNDP1 is an isoform of carnosinase that catalyzes the hydrolysis of dipeptides carnosine (β-alanyl-l-histidine) and homocarnosine (γ-aminobutyryl-lhistidine). Apart from amyloid-β, t-tau and p-tau, a decrease in CNDP1 was identified as a CSF biomarker for early-stage AD [46] and may serve a similar purpose in PDCI in our study. Exopeptidases like CPE/NFα1 and SCG3 are secretory sorting receptors that activate peptide messengers; target them at regulated secretory pathway yet get aggregated in senile plaques of AD patients and transgenic mice [47]. Hence, the down-regulation of *CPE* in our samples implies hints at possible aggregation. Dkk3 is a divergent glycoprotein co-expressed with amyloid-β peptide in plaques of AD patients [48]. It is enhanced in AD, DLB, and FTD. The mild reduction in our study suggests that Aβ plaques may not be involved in the pathogenesis of cognitive impairment in PD.

PD was associated with interesting proteins like *APP, NPTXR, CHI-3-L 1* and *SCG2*. APP binds to the death receptor DR6 via caspase [49] and its upregulation suggests axonal degeneration and neuronal death. *NPTXR*, [50] a transmembrane synaptic protein, is decreased in CSF of AD patients with severe cognitive impairment [51], hence the higher levels in our samples confirm normal cognition in PD. The significant down-regulation in *SCG3* in PDCI contrasted by an increase in *SCG2* in PD supports the hypothesis of catecholaminergic deficit in the later stages of PD [52, 53]; which is often riddled with cognitive impairment. *CHI3-L1* being associated with neuroinflammation [54] and faster cognitive decline, its increase in PD-CSF hints at the likely progress into cognitive impairment.

Although our primary aim was to identify biomarkers for PDCI, we serendipitously found higher fibrinogen levels in NPH, which tempts us to propose its likely contribution in the disease pathology. Other proteins of significance include ceruloplasmin, *IGFBP7, SERPINA1*, *VGF*-nerve growth factor, *NRCAM-L1* etc. Ceruloplasmin being an acute phase copper binding protein that lowers iron induced oxidative damage [55]; its mild reduction in NPH-CSF suggests iron deposition in the brain. Upregulation of *IGFBP7* in the hippocampus of AD mouse is associated with cognitive decline [56]; and partly explains the signs of cognitive deficits in our NPH patients. SerpinA1 is an anti-inflammatory protein of the choroid plexus, the reduction of which may have inflammatory consequences [29, 57]. Similar is the case with low VGF-nerve growth factor levels. Truncation of *NRCAM-L1* leads to hydrocephalus [58], hence its lower levels in our study may be a predictor of NPH, yet its pathogenic potential needs to be studied in detail. Although the sample size is small and is the limiting factor of our study, it provides convincing leads. Further, a study on a larger cohort and examination of plasma/serum would be essential to derive confirmatory answers.

## Conclusion

Our study suggests that *FGA* may be a better marker to distinguish PDCI from NPH. In view of the correlation between the digit-vigilance and N-BACK tests with CFAH and ApoA-IV levels respectively, these proteins may be useful in predicting cognitive impairment in PDCI and NPH. Since most proteins identified originate in the periphery, determining their levels in blood/serum may provide non-invasive avenues. Further validation in a larger cohort is essential to confirm the battery of proteins that can serve as biomarkers for PDCI. Thus, our non-targeted approach has provided essential clues for biomarker identification. Further studies using these proteins in animal models may assist the understanding of pathogenic trajectory of cognitive decline in PD.

## Supporting information

Proteomics MS supplementary.docx

## Abbreviations

ACN: Acetonitrile
AD: Alzheimer’s disease
AHSG: Fetuin-A
ANT: Animal naming test
ApoA-I: Apolipoprotein A1
ApoA-II: Apolipoprotein AII
ApoA-IV: Apolipoprotein-AIV
APP: Amyloid precursor protein
AVLT: Auditory verbal learning test
CFH/ CFAH: Complement factor H (Gene = *CFH*; protein = CFAH)
CFT: Complex figure test
CHI3L1: Chitinase-3-like protein 1
CLU: Clusterin
CNDP1: Carnosine dipeptidase I
COWA: Controlled Oral Word Association Test
CP: Ceruloplasmin
CPE: Carboxypeptidase E
CSF: Cerebrospinal fluid
CT: Color trails test
DKK3: Dickkpof 3
DLB: Dementia with Lewy bodies
DTT: Dithiothreitol
DWI: Diffusion weighted imaging
ELISA: Enzyme linked immunosorbent assay
FA: Formic acid
FASTA: Fast adaptive shrinkage thresholding algorithm
FBLN1: Fibulin 1
FGA: Fibrinogen α chain (Gene = *FGA*; protein = FIBA)
FGB: Fibrinogen β chain (Gene = *FGB*; protein = FIBB)
FGG: Fibrinogen γ chain (Gene = *FGG*; protein = FIBG)
FLAIR: Fluid attenuation inversion recovery
GSN: Gelsolin
HBA: Haemoglobin α chain
HBB: Haemoglobin β chain
HP: Haptoglobin
HRG: Histidine rich glycoprotein
IAA: Iodoacetamide
IGFBP7: Insulin-like growth factor-binding protein 7
IGHA2: Immunoglobulin heavy constant α 2
iTRAQ: Isobaric tags for relative and absolute quantitation
LC/MS-MS: Liquid chromatography mass spectrometry
LFQ: Label free quantitation
LRG1: Leucine-rich α-2 – glycoprotein
MDS: Movement disorders society
MOCA: Montreal cognitive assessment test
MRI: Magnetic resonance imaging
MSA: Multiple system atrophy
NCHL1: Neural cell adhesion molecule L1 like protein
NELL2: Neural EGFL like 2
NNC: Non-neurological control
NPH: Normal pressure hydrocephalus
NPTXR: Neuronal pentraxin receptor
NRCAM: Neuronal cell adhesion molecule
ORM1: Orosomucoid 1 or α-1 acid glycoprotein 1 precursor
ORM2: Orosomucoid 2 or α-1 acid glycoprotein 2 precursor
PDCI: PD with cognitive impairment
PDD: PD with dementia
PD-MCI: PD with mild cognitive impairment
Q-TOF: Quadrupole time of flight
SCG: Secretogranin
SCG2: Secretogranin-2 precursor
SCG3: Secretogranin 3 precursor
SERPINA1: α-1 antitrypsin
SERPINA3: α-1 anti-chymotrypsin
SERPING1: C1-inhibitor
SWI: Susceptibility weighted imaging
THY: Thy-1 membrane glycoprotein
TOL: Tower of London
UPDRS: Unified Parkinson’s disease rating scale
VGF: VGF Nerve growth factor
VTN: Vitronectin
WCST: Wisconsin card sorting test

## Acknowledgement

The authors are grateful to Indian Council for Medical Research, (BMS/TF/Trans-Neuro/2014-3424/Dec-15/48/MH/Govt) for funding support to PAA. AN was an ICMR-JRF [(No. 3/1/2/JRF-2014/HRD-102 (40832)]. Authors thank Mr. Darshak Gadara for help with Proteomics assays and Dr. Sathisha K for collating the mass spectrograms.

