## Supplementary material for "Fibrinogen and Complement Factor H are promising CSF protein biomarker(s) for Parkinson’s disease with cognitive impairment- A Proteomics and ELISA based study": Proteomics MS supplementary.docx

[Proteomics MS supplementary.docx](Proteomics%20MS%20supplementary.docx)

**Fig. S1**: Representative MRI images showing A. Brain of PDCI (yellow arrow) indicates the swallow tail on the left and its absence on the right. B. Brain of NPH (red arrow) indicates acute angle formed by the Corpus callosum, (white arrow) indicates crowded parasagittal sulci.

A.

B.

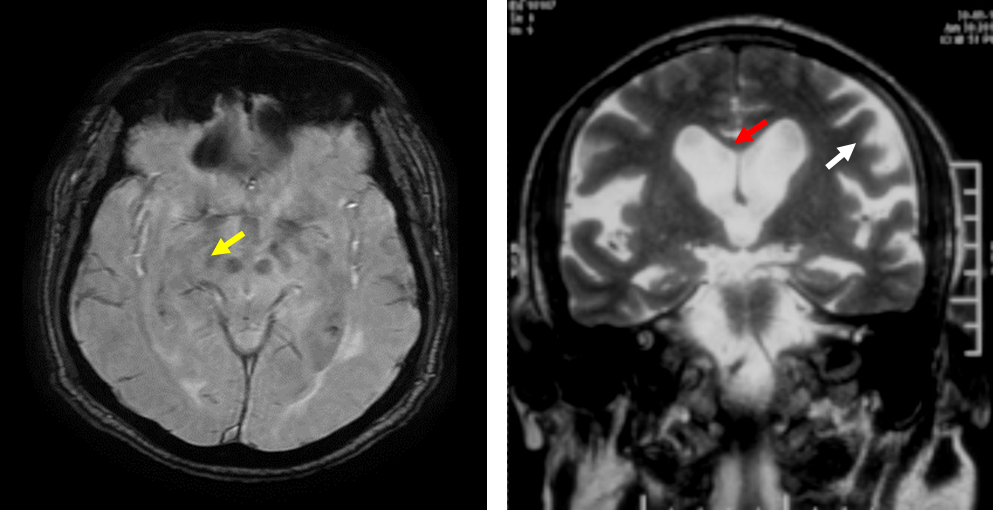

**Fig. S2**: Photomicrographs representing mass spectra of A. Fibrinogen α chain; B. Fibrinogen β chain; C. Fibrinogen γ chain; D. Gelsolin; E. Apo-AI; F. Apo-AIV

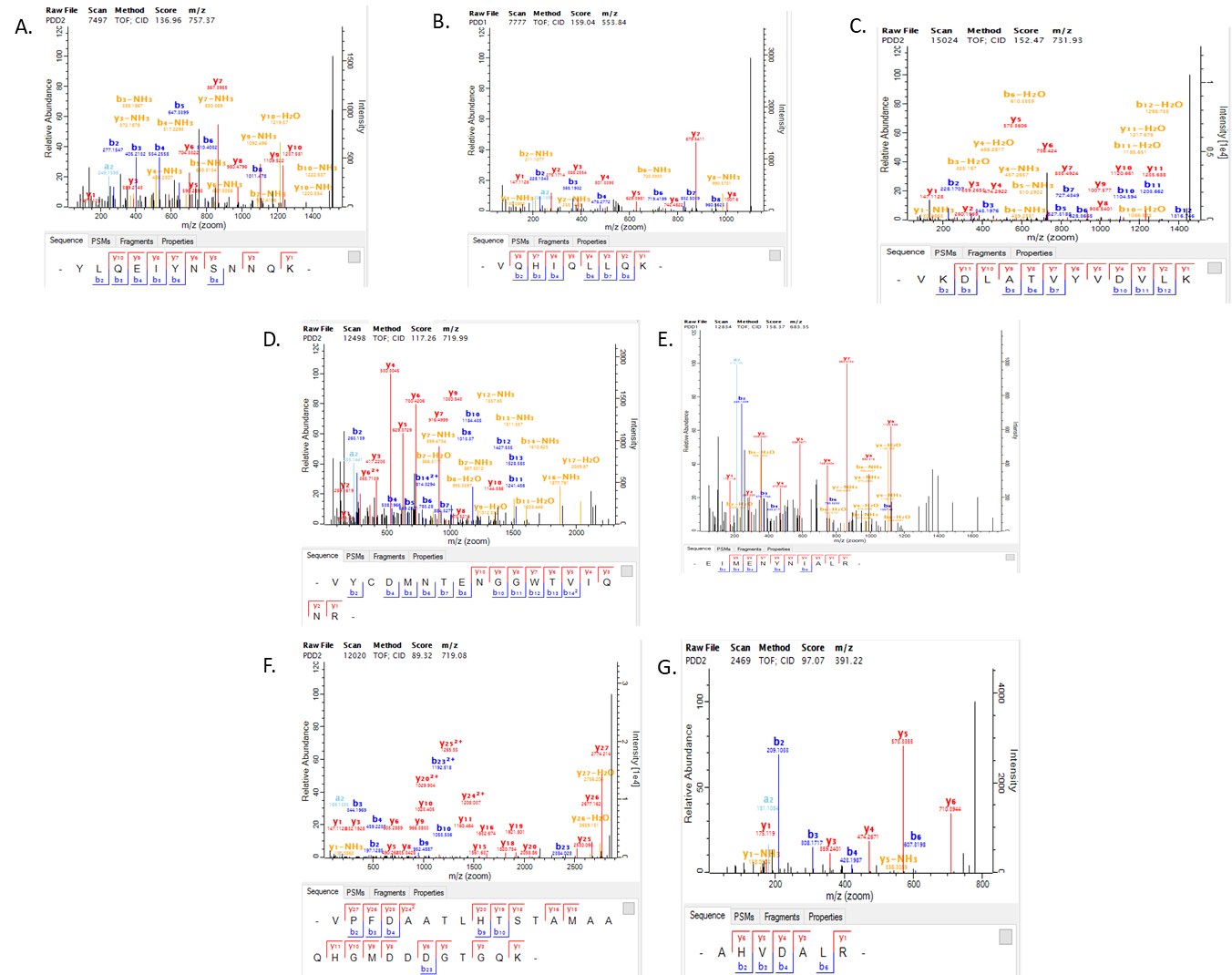

**Table S1:** Correlation between neuropsychological assessment test percentiles and ELISA based concentration of proteins viz. Fibrinogen, CFAH, APO-AI, APO-AIV, APO-AI:APO-AIV in NPH. Significant correlation coefficients and p-values are highlighted in red.

|  |  | Fibrinogen( conc ng/ml) | CFAH (conc pg/ml) | Apo-AI(conc ug/ml) | Apo-AIV concpg/ml | Apo-AI:Apo-AIV |
| --- | --- | --- | --- | --- | --- | --- |
| MOCA | Correlation Coefficient | -.138 | -.092 | .506^*^ | -.255 | .036 |
|  | Sig. (2-tailed) | .587 | .717 | .046 | .383 | .901 |
|  | N | 10 | 10 | 10 | 8 | 8 |
| COWA percentile | Correlation Coefficient | -.500 | -.567^*^ | -.167 | -.619^*^ | .371 |
|  | Sig. (2-tailed) | .082 | .049 | .562 | .044 | .228 |
|  | N | 9 | 9 | 9 | 8 | 8 |
| ANT percentile | Correlation Coefficient | -.514 | -.400 | -.400 | -.618^*^ | .400 |
|  | Sig. (2-tailed) | .058 | .140 | .140 | .034 | .170 |
|  | N | 9 | 9 | 9 | 8 | 8 |
| Verbal N back 2 ERROR percentile | Correlation Coefficient | -.057 | 0.000 | .229 | -.741^*^ | .519 |
|  | Sig. (2-tailed) | .833 | 1.000 | .399 | .012 | .079 |
|  | N | 9 | 9 | 9 | 8 | 8 |
| AVLT IR percentile | Correlation Coefficient | 0.000 | .065 | .196 | -.732^*^ | .394 |
|  | Sig. (2-tailed) | 1.000 | .818 | .489 | .029 | .241 |
|  | N | 9 | 9 | 9 | 7 | 7 |
| WCST no. of categories percentile | Correlation Coefficient | -.403 | -.645^*^ | -.161 | -.053 | -.265 |
|  | Sig. (2-tailed) | .189 | .036 | .599 | .874 | .427 |
|  | N | 8 | 8 | 8 | 7 | 7 |

**Table S2:**

|  |  | Fibrinogen( conc ng/ml) | CFAH (conc pg/ml) | Apo-AI(conc ug/ml) | Apo-AIV concpg/ml | Apo-AI:Apo-AIV |
| --- | --- | --- | --- | --- | --- | --- |
| WCST no. of categories percentile | Correlation Coefficient | .414 | .276 | .138 | .966^**^ | -.966^**^ |
|  | Sig. (2-tailed) | .251 | .444 | .702 | .007 | .007 |
|  | N | 6 | 6 | 6 | 6 | 6 |
